# Identification of immune-related genes for the diagnosis of ischaemic heart failure based on bioinformatics and three machine learning models

**DOI:** 10.1101/2023.05.22.541684

**Authors:** Yi-ding Yu, Yi-tao Xue, Yan Li

**Author notes:** **Correspondence:** Correspondence should be addressed to Yi-tao Xue; and Yan Li;.

## Abstract

**Background:** The role of immune cells in the pathogenesis of ischaemic heart failure (IHF) is well-established. However, identifying key diagnostic candidate genes in patients with IHF remains a challenge. Therefore, this study aimed to use bioinformatics and machine learning algorithms to identify potential diagnostic genes for IHF.

**Methods:** Two IHF datasets were obtained from the GEO database, and key genes for IHF were identified using Limma and WGCNA. Functional enrichment analysis was performed to explore the potential mechanisms of IHF. Next, we used three machine learning algorithms, namely LASSO, RF, and SVM-REF, to identify immune-related diagnostic genes for IHF. ssGSEA enrichment analysis was also completed. To assess the diagnostic value of the identified genes, we developed nomogram and validated them on additional GEO datasets. Finally, an immune infiltration analysis was conducted using the CIBERSORT algorithm to explore immune cell dysregulation in IHF.

**Results:** Our analysis yielded a total of 92 key genes associated with IHF. Enrichment analysis revealed that the mechanisms underlying IHF are mostly associated with immunity and inflammation. Using the machine learning algorithms, we identified four IHF diagnostic genes, namely RNASE2, MFAP4, CHRDL1, and KCNN3. We constructed nomogram and validated the diagnostic value of these genes on additional GEO datasets. The results showed that these four genes had high diagnostic value (AUC value of 0.961). Furthermore, our immune infiltration analysis revealed the presence of three dysregulated immune cells in IHF, namely Macrophages M2, Monocytes, and T cells gamma delta.

**Conclusion:** In summary, our study identified four potential diagnostic candidate genes for IHF by using bioinformatics and machine learning algorithms. We also explored the potential molecular mechanisms of IHF and the immune cell infiltration environment of the failing myocardium. These findings provide new insights into the pathogenesis, diagnosis, and treatment of IHF.

## 1. INTRODUCTION

Cardiovascular disease is a leading cause of human mortality, encompassing hypertension, arrhythmias, coronary heart disease, and heart failure, among other heart-related conditions. Of these, heart failure is often the ultimate outcome of most cardiovascular diseases, primarily due to structural changes or functional impairments of the heart that hinder ventricular filling or ejection function^1^. With the advent of interventional techniques and drugs, survival rates for patients with ischemic heart disease and acute myocardial infarction are improving, leading to an increased number of patients at risk for heart failure^2^. Damaged and unrecoverable myocardium is the principal cause of heart failure and death in patients within five years of an infarction^3^. Consequently, identifying myocardial-specific genes for ischemic heart failure (IHF) at the transcriptome level could facilitate early diagnosis and targeted treatment of these patients.

Previous studies have demonstrated that immune cells play protective and destructive roles in cardiac remodeling after infarction^4^. Multipotent cells of the innate immune system, such as monocytes and macrophages, are critical for the initial inflammatory response to post-infarction myocardial injury and subsequent wound healing^5^. Clinical studies have aimed to target elements of the immune response in heart failure by modulating the inflammatory response. However, the results of these clinical studies have been unsatisfactory, and in some cases, have even exacerbated heart failure^6^. Therefore, examining the immune response and associated genes in the progression of IHF could help develop novel therapeutic strategies.

Rapid advances in high-throughput technologies and bioinformatics can aid in the screening of sensitive and specific diagnostic tools to diagnose and treat heart failure patients before they reach a refractory end stage. Additionally, with the development and maturation of machine learning in bioinformatics applications, multiple machine learning models can aid in uncovering potential mechanisms, prospective diagnostic tools, and therapeutic targets for IHF^7^.

In this study, we initiated by acquiring two IHF datasets from the GEO database, followed by identifying differentially expressed genes (DEGs) through the Limma package. Subsequently, we selected significant modular genes via weighted gene co-expression network analysis (WGCNA). We then performed functional enrichment analysis and constructed a protein-protein interaction network (PPI). We utilized three different machine learning models, namely, least absolute shrinkage and selection operator (LASSO), random forest (RF), and support vector machine recursive feature elimination (SVM-REF), to determine diagnostic models. We further assessed these genes through nomogram and ssGSEA enrichment analysis. Additionally, we validated the diagnostic models on another GEO dataset. Finally, we conducted immune cell infiltration analysis on the dataset to identify immune-related genes in IHF and to elucidate the role of immune cells in the development of this condition.

## 2. MATERIALS AND METHODS

### 2.1 Microarray Data

Figure 1 depicts the study flowchart. We downloaded GSE21610 and GSE76701 from the GEO database for the GPL570 platform as the IHF dataset, and GSE57338 as the validation set^8–11^.

**Figure 1:**
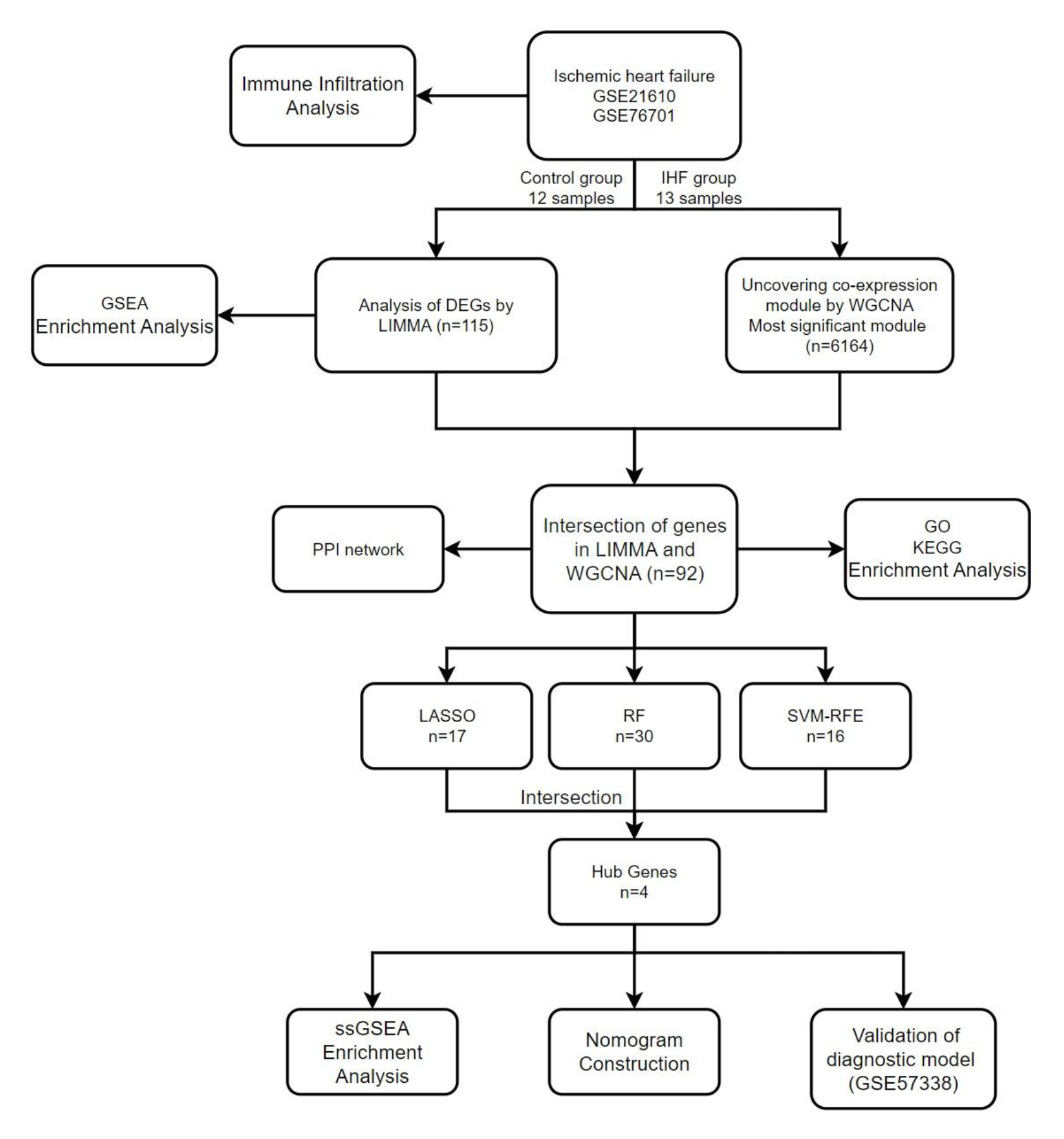
*The study flowchart*.

### 2.2 Data Processing and Differentially Expressed Gene Screening

To collate and analyze our data, we utilized R software (version 4.2.0) and accessed the GEO database through the GEOquery package to download the GSE21610 and GSE76701 datasets. To ensure accuracy and consistency, we removed probes corresponding to multiple molecules and retained only the probe with the highest signal value for each molecule. We utilized the ComBat function of the sva package to eliminate batch effect from the filtered data. To gain insights into the differences between heart failure and normal samples, we employed the limma package and identified genes with p-values < 0.05 and | log2(FC)| ≥ 1 as the difference genes. We further visualized our findings using the ggplot2 package and the pheatmap package^12–15^. To assess normalization, we utilized a box plot, and for clustering between sample subgroups, we used a PCA plot.

### 2.3 Weighted Gene Co-Expression Network Analysis and Module Gene Selection

In addition to obtaining differential genes with the limma package, we also used the WGCNA package to explore gene modules with relatedness^16^. A scale-free co-expression network was created after removing ineligible genes and samples by the goodSamplesGenes function with a filtering criterion of 0.5. Subsequently, adjacency was calculated by default using β = 30 and scale-free R2 = 0.9 as a soft threshold, and adjacency was converted to a topological overlap matrix (TOM), which was used to determine gene ratios and dissimilarity. Genes with the same expression profile were grouped into gene modules using average linkage hierarchical clustering. We preferred larger modules, so we set the minimum module size to 200. Finally, we calculate the similarity of the modules’ signature genes, select the cut lines of the module dendrogram to combine several modules for the next step of the study, and complete the visualisation of the signature gene network. WGCNA analysis was used to identify important modules in IHF.

### 2.4 Functional Enrichment Analysis

We utilized the clusterProfiler package to conduct Gene Ontology (GO), Kyoto Encyclopedia of Genes and Genomes (KEGG), and Gene Set Enrichment Analysis (GSEA) enrichment analyses and visualizations^17–20^. Specifically, we performed GSEA enrichment analysis on all genes within our dataset. Subsequently, we identified the key genes of IHF by intersecting the DEGs with the important module genes of WGCNA. Finally, we performed GO and KEGG enrichment analyses on the key genes of IHF.

### 2.5 Protein–Protein Interaction Network Construction

To mine the interactions between protein-coding genes, we built a PPI network using the STRING database^21^. We uploaded the list of IHF key genes to the STRING database with the following assay settings: Network type: full STRING network, meaning of network edges: evidence, minimum required interaction score. medium confidence (0.400). We then imported the results into Cytoscape 3.6.1 for visualisation and subsequent analysis^22^.

### 2.6 Machine Learning

We will use three machine learning algorithms, LASSO, RF and SVM-RFE, to further screen candidate genes for IHF diagnosis^23–25^. We use the glmnet package to perform the LASSO algorithm, choosing ten-fold cross-validation to select the prominent genes. We used the randomForest package to complete the RF algorithm, selecting the top 30 genes as alternative genes. We used the e1071 package to complete the SVM-RFE algorithm to select the number of genes with the highest precision as candidate genes. After completing the calculation, we selected the intersection of the three as the diagnostic genes for IHF.

### 2.7 Nomogram Construction and Validation of diagnostic model

We utilized the rms package to develop a nomogram for the identification of IHF diagnostic genes^26^. Points represent the scores of candidate genes and Total Points represent the sum of all the above gene scores. Subsequently, box plots of gene expression were generated and receiver operating characteristic (ROC) curves were constructed to determine the diagnostic value of the candidate genes. The area under the curve (AUC) was calculated to quantify their value, with an AUC value greater than 0.7 being considered as the ideal diagnostic threshold. To further validate our findings, we conducted an analysis of individual and combined genes using the GSE57338 dataset. We assessed the discriminatory ability of the diagnostic model by ROC curves once again.

### 2.8 ssGSEA Enrichment Analysis

We used the clusterProfiler package to complete ssGSEA enrichment analysis of genes from the IHF diagnostic model to explore the function of these genes in the IHF process^27^.

### 2.9 Immune Infiltration Analysis

We used the CIBERSORT package to assess the content of immune cells and stromal cells in IHF myocardial samples to depict a cellular heterogeneous landscape of myocardial expression profiles and to complete the immune cell infiltration analysis^28^. Bar charts were used to visualise the proportion of each type of immune cell in the different samples. Differences in cell distribution between the IHF and normal groups were compared by t-test, with the cut-off value set at p < 0.05.

## 3. RESULTS

### 3.1 Identification of Differentially Expressed Genes

After data merging and normalisation, we identified a total of 115 differential genes in the merged dataset using the LIMMA package, of which there were 78 up-regulated and 37 down-regulated genes. The visualisation of the associated results was shown in Figure 2.

**Figure 2:**
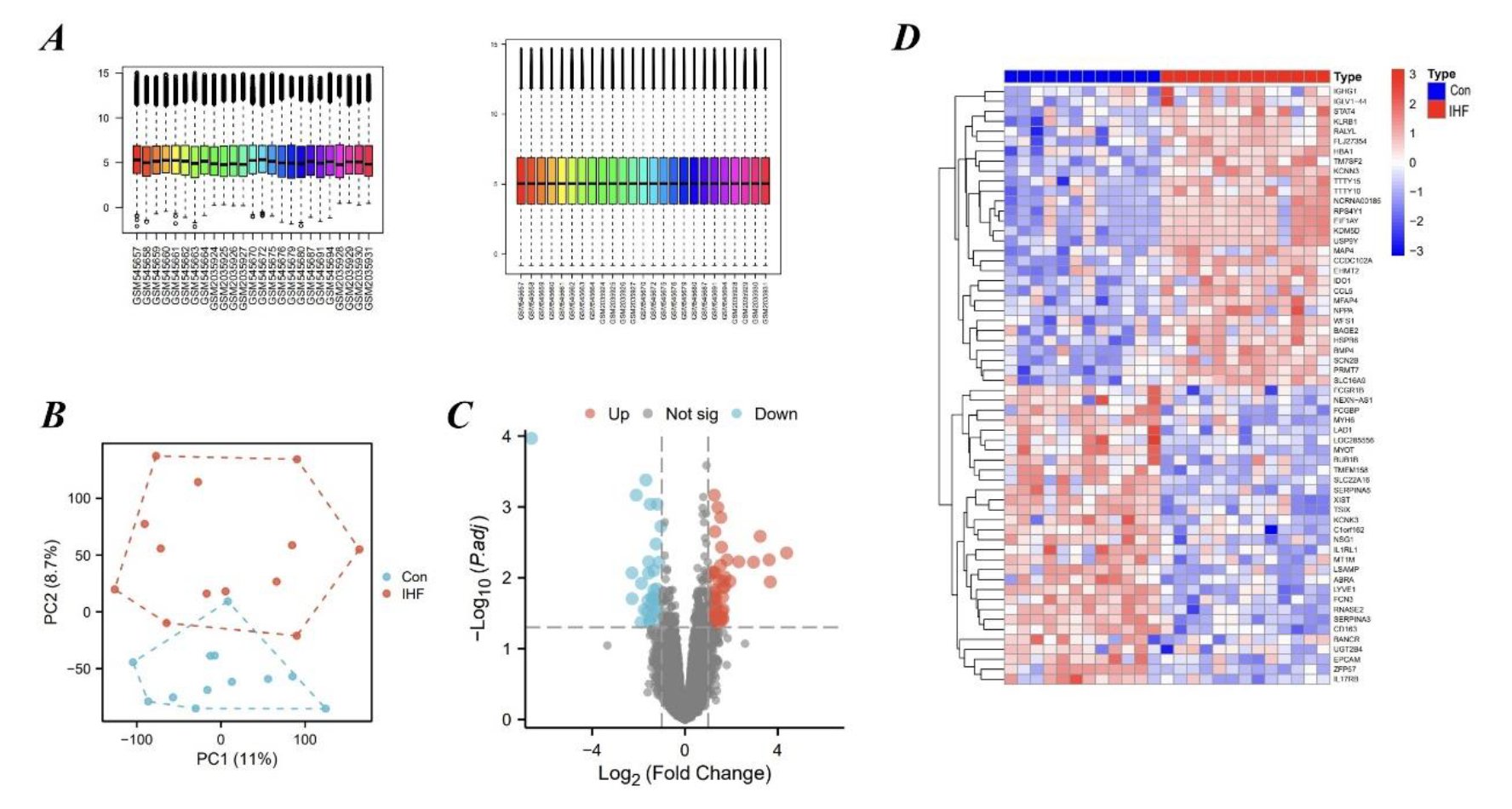
*Results of Differentially Expressed Genes. (A) Normalization of data. (B) Principal Component Analysis plot. (C) Volcano plot. (D) Heatmap plot*.

### 3.2 Weighted Gene Co-Expression Network Analysis and Key Module Identification

We used WGCNA to identify the most relevant modules in IHF. Ultimately, we obtained 16 gene co-expression modules. Among them, the turquoise module had the highest correlation with IHF (correlation coefficient = 0.72, p = 5e-05), containing a total of 6164 genes. Therefore, we selected the turquoise module as the key module for the subsequent analysis. The visualization of the results was shown in Figure 3.

**Figure 3:**
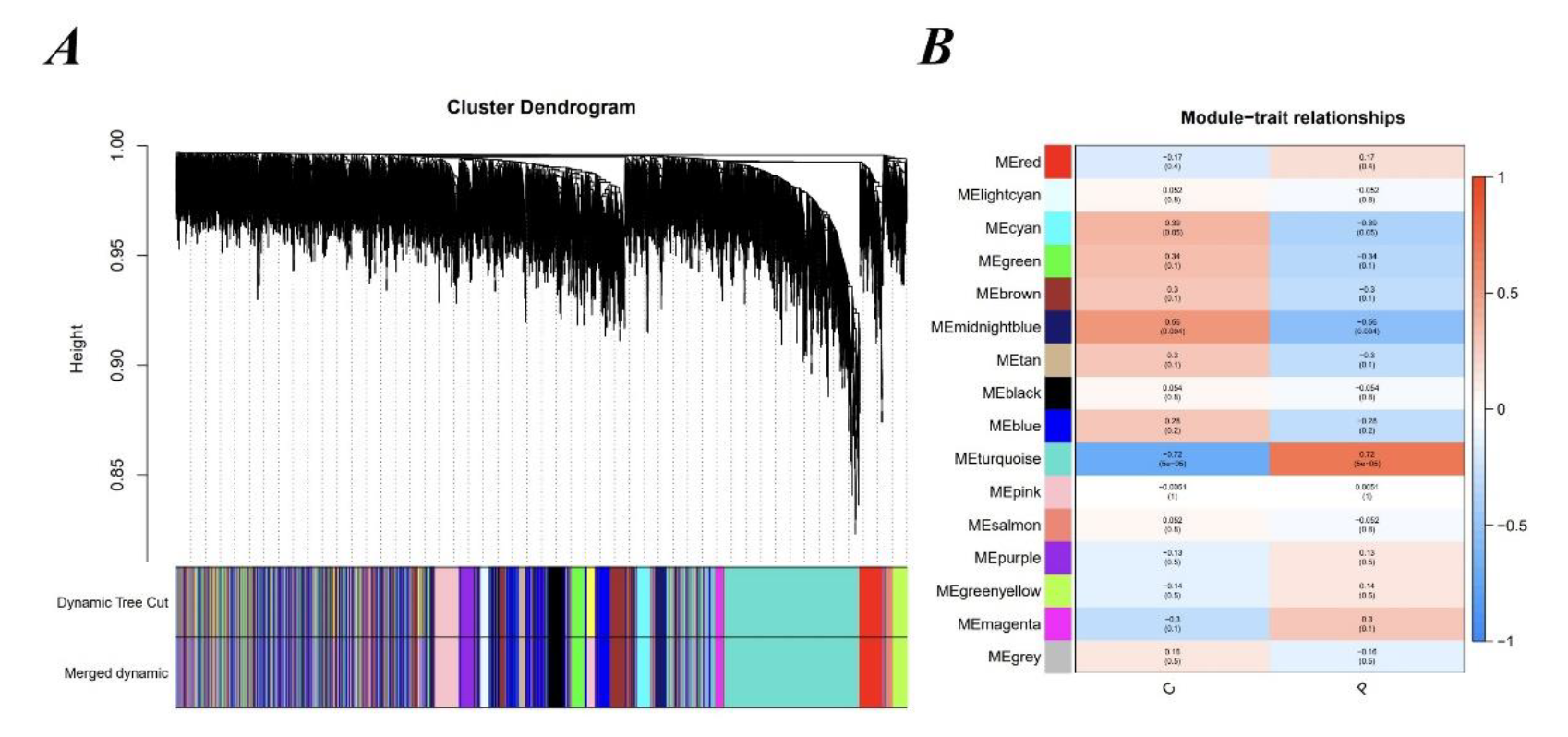
*WGCNA analysis result. (A) Gene co-expression modules represented by different colors under the gene tree. (B) 16 gene co-expression modules*.

### 3.3 Functional Enrichment Analysis of Ischemic Heart Failure

To ensure that genes with significant functions but insignificant differential multiplicities were not overlooked, we conducted a GSEA enrichment analysis on all the genes in the dataset. The GSEA results indicated that the genes likely to be associated with IHF were Antigen processing and presentation, Apoptosis, Proteasome, and Th1 and Th2 cell differentiation, which are involved in biological functions such as programmed cell death, inflammation and immunity. Afterwards, we intersected DEGs with the turquoise module genes to obtain 92 IHF key genes. We performed GO and KEGG enrichment analysis on the IHF key genes. the categories of GO analysis include biological process (BP), cellular component (CC) and molecular function (MF), and we will show the top five results respectively. The KEGG enrichment analysis did not yield many pathways, so we will show all the enriched results. The visualisation results were shown in Figure 4.

**Figure 4:**
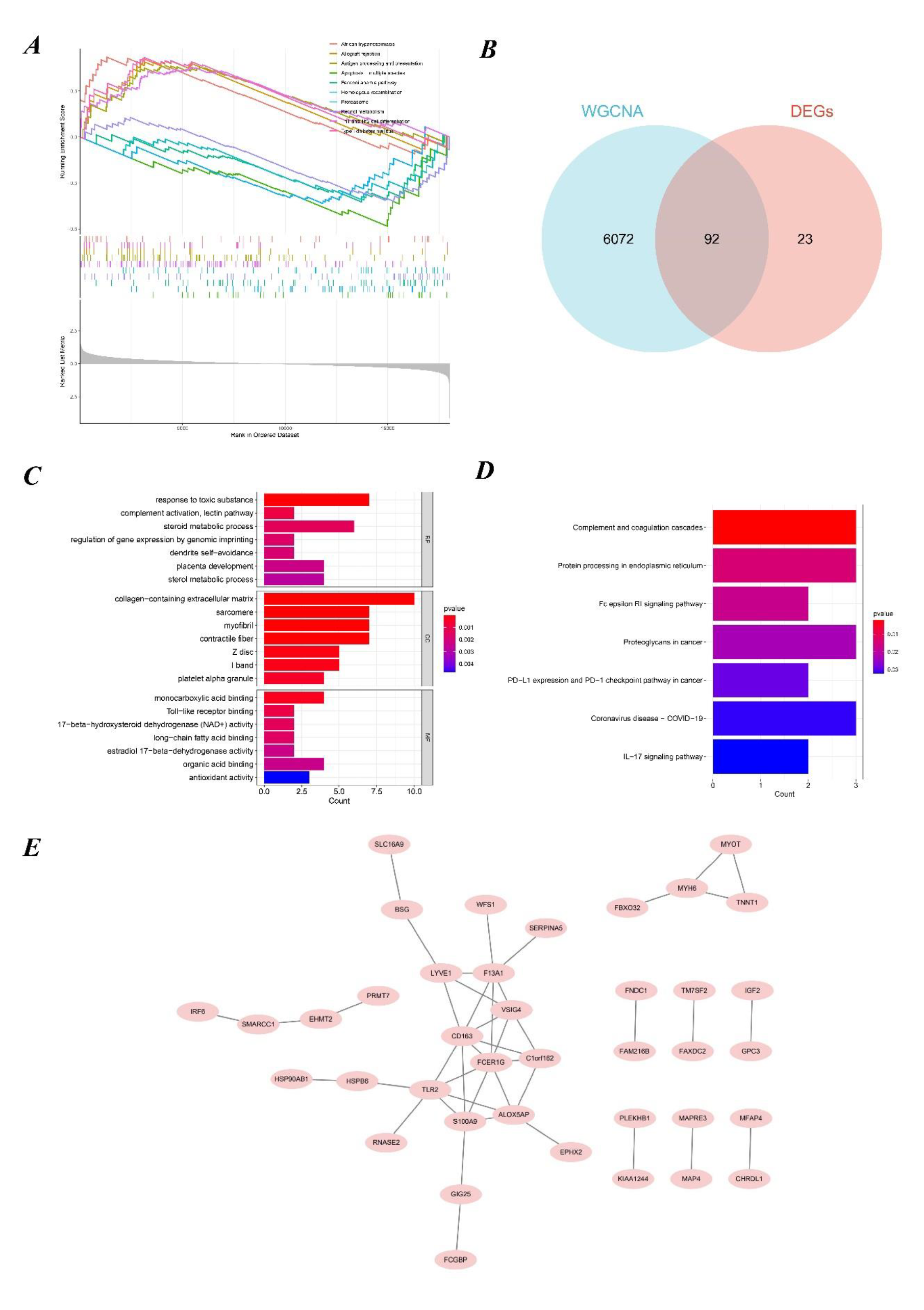
*Functional Enrichment Analysis of IHF and PPI network. (A) GSEA enrichment analysis results. (B) The intersection of DEGs and the turquoise module. (C) GO enrichment analysis results. (D) KEGG enrichment analysis results. (E) PPI network*.

### 3.4 PPI network construction

We entered the IHF key genes into the STRING database for PPI network construction, which resulted in the identification of 39 nodes and 44 interactions. The visualisation result was shown in Figure 4-E.

### 3.4 Identification of Hub Genes via Machine Learning

We used three machine learning algorithms, LASSO, RF and SVM-RFE, to further screen Hub genes for IHF. We identified 17 potential candidate biomarkers by the LASSO algorithm. the RF algorithm ranked the genes based on the importance calculation of each gene, and we selected the top 30 as potential candidates for IHF. The SVM-RFE algorithm exhibited the highest precision, identifying 13 genes with a constant precision score of 1 thereafter. To establish the optimal number of Hub genes, we selected the top 16 genes for the SVM-RFE algorithm results as candidate genes. By intersecting the results of all three algorithms, we identified four Hub genes for IHF: RNASE2, MFAP4, CHRDL1, and KCNN3. The visualisation results were shown in Figure 5.

**Figure 5:**
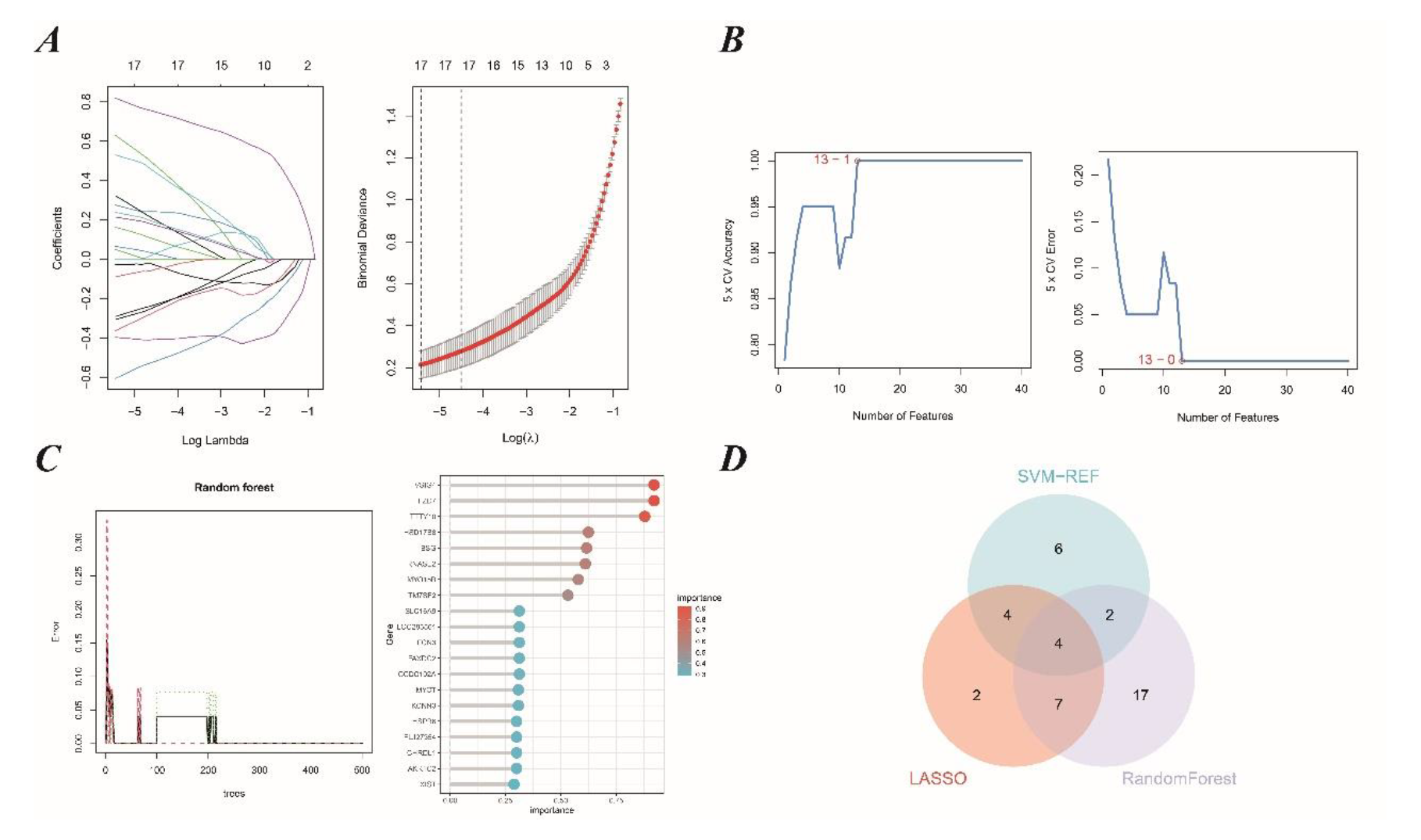
*Machine learning in screening candidate diagnostic biomarkers for IHF. (A) Biomarkers screening in the Lasso model. (B)Biomarkers screening in the SVM-RFE model. (C) Biomarkers screening in the RF model. (D) Venn diagram shows that four candidate diagnostic genes are identified via the above three algorithms*.

### 3.5 Diagnostic Value Assessment

We constructed nomograms based on the four Hub genes and built ROC curves to assess the diagnostic specificity and sensitivity of each gene and nomogram. In addition, we plotted differential expression box plots for the Hub genes. Finally, we completed the validation of Hub genes in GSE57338 using ROC curve analysis. The validation results showed that the AUC of each gene was greater than 0.7, and the AUC of the 4-gene diagnostic model was 0.961, which had high diagnostic value. The visualisation results were shown in Figure 6.

**Figure 6:**
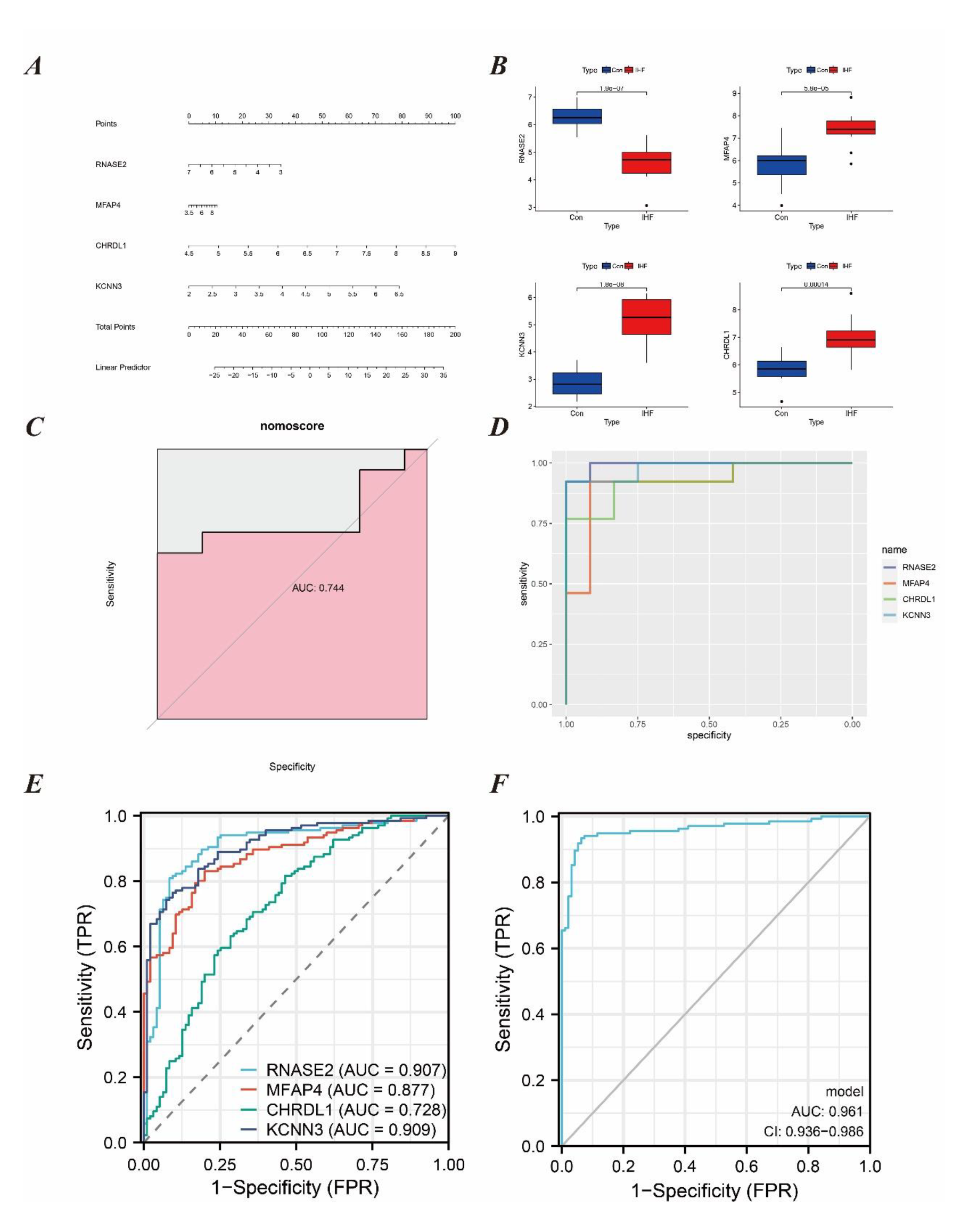
*Results of Diagnostic Value Assessment. (A) The visible nomogram for diagnosing IHF. (B) Differential expression box plots for the Hub genes. (C) The ROC curve of nomogram showed the IHF diagnostic value. (D) The ROC curve of each candidate genes showed the IHF diagnostic value. (E) The ROC curve of each candidate genes in GSE57338. (F) The ROC curve of the 4-gene diagnostic model in GSE57338*.

### 3.6 ssGSEA Enrichment Analysis

We performed ssGSEA enrichment analysis for RNASE2, MFAP4, CHRDL1, and KCNN3, respectively. The results showed that all these genes are involved in the development of IHF and the biological functions of immunity and inflammation in the disease process to varying degrees. For example, RNASE2 is involved in Adrenergic signaling in cardiomyocytes, Cell cycle, Cellular senescence, IL-17 signaling pathway, and Proteasome. MFAP4 is involved in TKCNN3 is involved in the TGF-beta signaling pathway. CHRDL1 is involved in the Proteasome, Primary immunodeficiency and Circadian CHRDL1 is involved in Proteasome, Primary immunodeficiency and Circadian rhythm. Detailed results and visualisation were shown in Figure 7.

**Figure 7:**
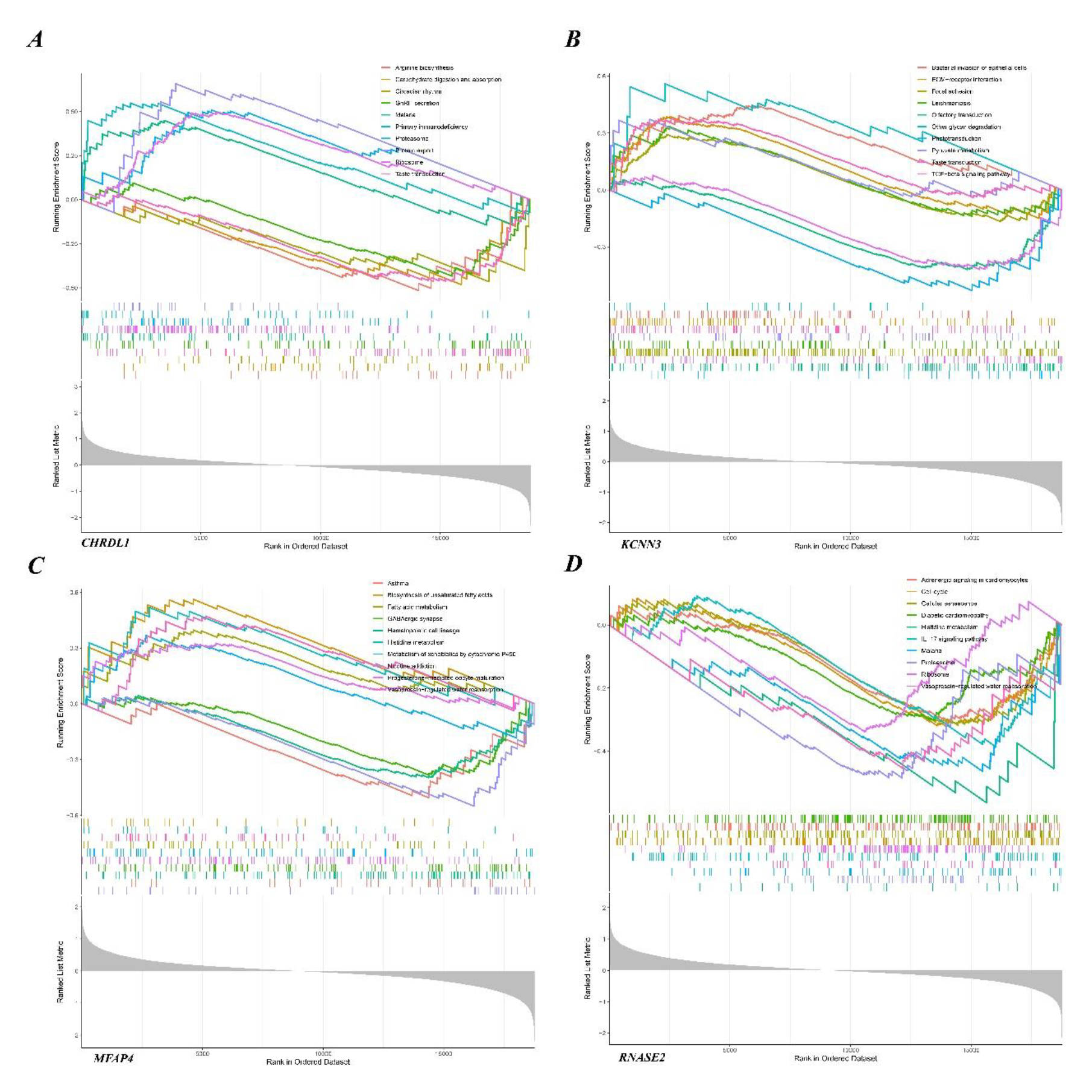
*The results of ssGSEA Enrichment Analysis. (A) The results of CHRDL1. (B) The results of KCNN3. (C) The results of MFAP4. (D) The results of RNASE2*.

### 3.7 Immune Cell Infiltration Analysis

Due to the important role of immune cells in the development of IHF, we also performed an immune infiltration analysis of the dataset by means of the CIBERSORT algorithm. The bar charts clearly show the content of the different subpopulations in each sample. We assessed the heterogeneity of cell composition between the heart failure samples and the healthy samples and the results showed that there were three immune cell infiltrates that were significantly different. Macrophages M2 and Monocytes were higher in normal samples than in heart failure samples, and T cells gamma delta was lower than in heart failure samples. The differential infiltration of these 3 types of immune cells may provide potential regulatory points for the treatment of IHF. The visualisation results were shown in Figure 8.

**Figure 8:**
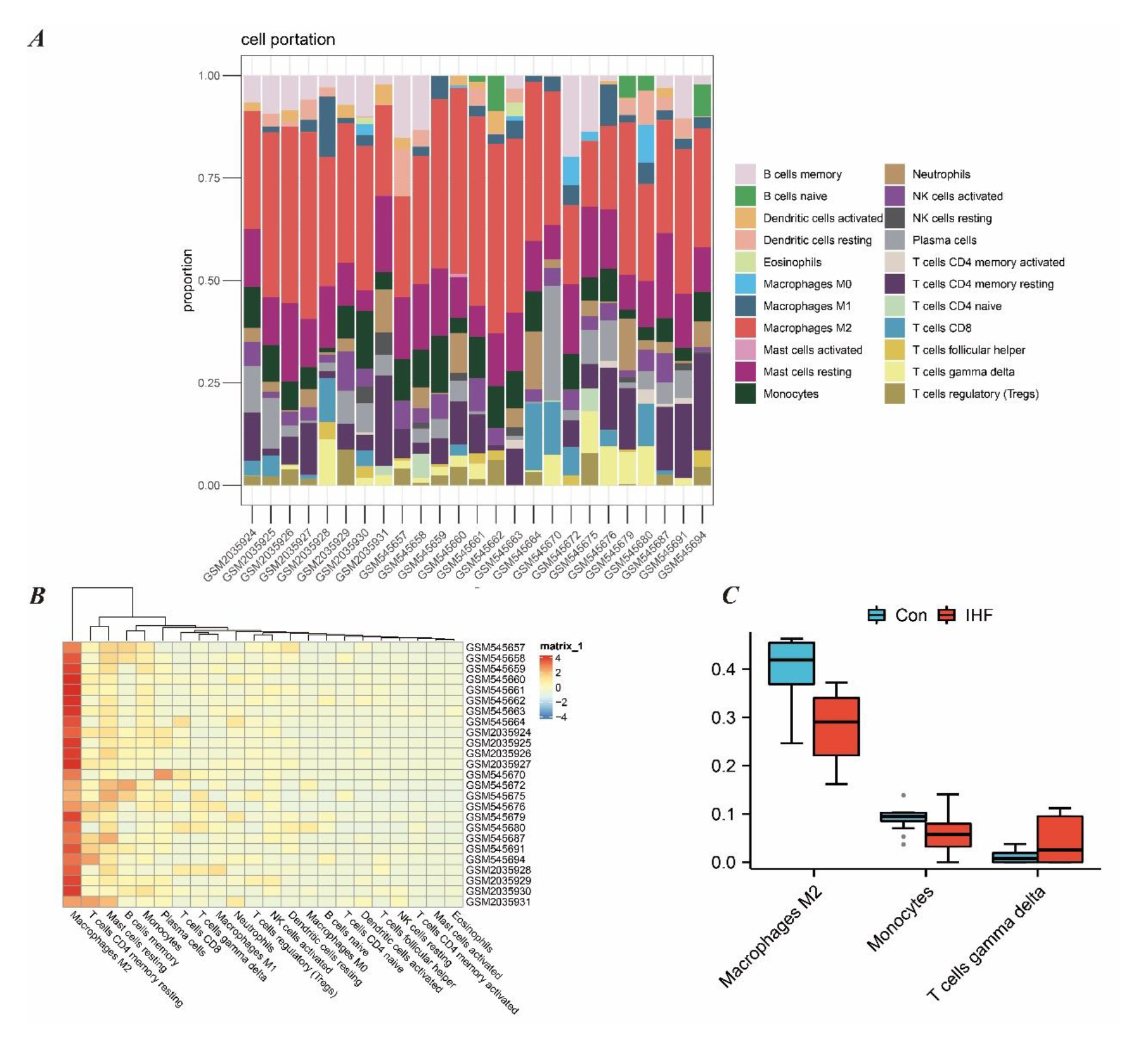
*Immune cell infiltration analysis between IHF and control. (A) The proportion of 22 kinds of immune cells in different samples visualized from the bar plot. (B) The expression of the 22 immune cells in the different samples can be seen in the heat map. (C) Expression of the 3 dysregulated immune cells in the IHF and controls as seen in the box plot*.

## 4. DISCUSSION

HF is always a major public health challenge. Among them, left ventricular systolic dysfunction due to obstructive coronary artery disease is the most common cause of heart failure worldwide. While new treatments such as mechanical unloading and modulation of the inflammatory response look promising, understanding the mechanisms of IHF is a key step in finding new ways to address the risk of heart failure^29^.

In this study, we explored the potential mechanisms of IHF through three enrichment analyses. The results suggest that the underlying mechanisms of IHF are mostly related to inflammation, immunity, and metabolism. For example, Antigen processing and presentation and Th1 and Th2 cell differentiation in the enrichment results are involved in cellular immune processes. It has been demonstrated that cardiac fibroblasts take up and process antigens and promote cardiac fibrosis and dysfunction through IFNγ^30^. Inflammatory cytokines such as IFNγ, TNF-α, IL-2, and IL-6 secreted by Th1 and Th2 cells have also been important players in the development of heart failure^31, 32^. The Proteasome involves the ubiquitin-proteasome system in maintaining protein homeostasis and cardiac^33^. The ubiquitin-proteasome system plays a key role in maintaining protein homeostasis and cardiac function. On the one hand, the ubiquitin-proteasome system is the major protein degradation system involved in the regulation of inflammation and selective mitochondrial autophagy during heart failure^34^. On the other hand, this system is also involved in the development of heart failure in terms of cardiac energy metabolism^35^. In addition, how to mitigate the cardiotoxicity induced by proteasome inhibitors of anticancer drugs has been a popular research in recent years^36^. We also used bioinformatics tools and machine learning algorithms to screen for Hub genes for IHF, namely RNASE2, MFAP4, CHRDL1, and KCNN3, and validated them in a larger dataset. The AUC value of this 4-gene diagnostic model in the validation set was 0.961, implying that this diagnostic model has a high diagnostic value. Nomograms were also plotted to aid the application of this diagnostic model. We completed the ssGSEA analysis of these 4 genes in order to provide new ideas for our understanding of the molecular mechanism of IHF.

Ribonuclease A family member 2 (RNASE2) is a non-secretory ribonuclease that belongs to the RNaseA superfamily. Also known as an eosinophil-derived neurotoxin (EDN), RNASE2 is involved in immune and inflammatory related pathways^37^. Its broad antiviral activity is primarily directed against single-stranded RNA viruses, such as human immunodeficiency virus^38^. Studies have shown that RNASE2 acts as a catalyst for human dendritic cells and promotes the secretion of a plethora of cytokines and chemokines^39^. RNASE2 also functions as an endogenous ligand for toll-like receptor (TLR) 2. On the one hand, downstream pathway stimulation via TLR may activate myeloid differentiation factor 88 (MyD88) and mitogen-activated protein kinase (MAPK) to promote the production of pro-inflammatory factors such as IL- 10^40, 41^. On the other hand, TLR immunosensing against live pathogens may also allow RNASE2 to act as a bridge between innate and adaptive immunity^42^. In the cardiovascular context, TLR2 has been reported to regulate myocardial ischaemia, and thus RNASE2 may be involved in innate immune responses in the pathogenesis of IHF via TLR2^43^.

Microfibrillar Associated Protein 4 (MFAP4) is an extracellular matrix protein that belongs to the fibrinogen-associated protein superfamily. Vascular smooth muscle cells produce MFAP4, which is highly enriched in the vessels of the heart and lungs, and is believed to contribute to the structure and function of elastic fibers^44^. Thus, MFAP4 holds significant research value in arterial vascular-related diseases. Studies have shown that MFAP4 can induce the proliferation and migration of vascular smooth muscle cells and promote monocyte chemotaxis, which can accelerate neointimal proliferation after vascular injury^45^. Additionally, two clinical studies have demonstrated the potential of MFAP4 as a biomarker of atherosclerotic disease. In patients with stable coronary artery disease, serum MFAP4 levels were lower compared to patients with acute infarction^46^. Moreover, MFAP4 has the potential to serve as a biomarker for assessing the degree of coronary artery stenosis in acute heart attacks^47^. Concerning heart failure, MFAP4 plays a crucial role in macrophage infiltration, inflammation, and myocardial fibrosis^48^. MFAP4 knockout experiments have shown that MFAP4 deficiency can lead to dysregulated integration of G protein-coupled receptors and integrin signaling in the heart, which exacerbates cardiomyocyte hypertrophy^49^.

Chordin-like 1 (CHRDL1) is a secreted protein that acts as an antagonist of bone morphogenetic protein (BMP)^50^. There has been little research on the association of CHRDL1 with heart disease, but recently Mauro Giacca and his team have identified the cardioprotective effects of CHRDL1 by a method called cardiac FunSel^51^. Its cardioprotective effect stems from the maintenance of cardiomyocyte viability by blocking the negative effect of BMP4 on cardiomyocyte autophagy. Also, CHRDL1 inhibits post-infarction cardiac fibrosis and enables post-infarction remodelling by inhibiting the negative effects of TGF-β on cardiac fibroblasts. In addition, a chromosomal genetic analysis showed that CHRDL1 was strongly associated with lowered lipids, suggesting that CHRDL1 has potential in coronary plaque control and may improve myocardial supply and treat IHF by lowering LDL and reversing plaque^52^. Potassium calcium-activated channel subfamily N member 3(KCNN3) belongs to the KCNN family of potassium channels. It encodes an integral membrane protein that forms a voltage-independent calcium-activated channel, which is thought to regulate neuronal excitability by contributing to the slow component of synaptic AHP^53^. Thus, KCNN3 has potential in the treatment of arrhythmias, especially since GWAS evidence suggests that variants in the KCNN3 gene are associated with atrial fibrillation^54^. In the field of heart failure, it has been shown that ventricular expression of KCNN3 is significantly increased in heart failure when ventricular tissue from patients with heart failure is compared to healthy samples, in line with our findings. It has also been shown that upregulation of KCNN3 leads to deterioration of ventricular function, which may be related to the involvement of this gene in the induction of ventricular tachycardia, but the specific mechanisms behind this need further investigation^56^.

The results of the immune infiltration analysis showed that there were three types of immune cell infiltration that were significantly different. Macrophages were divided into two subpopulations based on their function and level of inflammatory factor secretion: Macrophages M1 and Macrophages M2. Macrophages M2 have an anti-inflammatory effect and are mainly activated by IL-4 inflammatory factor, inhibiting M1 macrophages mainly by secreting anti-inflammatory cytokines such as IL-10, which play a role in processes such as wound healing and tissue repair^57^. Macrophages M2 are lower in heart failure samples than in normal samples, and imbalanced M1/M2 macrophages may exacerbate inflammatory damage to cardiomyocytes, exacerbating myocardial dysfunction and fibrosis^58^. Monocytes play a key role in orchestrating the inflammatory cascade response and the pathogenesis of HF. However, the highly differentiated nature of Monocytes complicates their function. On the one hand, Monocytes are one of the major cellular targets of pro-inflammatory cytokines, and TNF-α induces monocytes to promote NO synthase production, thereby inducing apoptosis in cardiomyocytes, which in turn leads to further activation of the cascade by Monocytes, resulting in a vicious cycle in the failing myocardium^59^. On the other hand, it has been suggested that activated monocytes can infiltrate the myocardium in order to exert phagocytic and reparative effects^60^. This may explain the general lack of clinical success in treating HF by modulating the cytokine system. The main biological effect of T cells gamma delta (Tgd cells) is cytotoxicity, which may account for its elevation in heart failure samples. Studies have shown that modulation of the IL- 17A/Tgd cells axis can effectively modulate inflammation levels and slow down the process of myocardial fibrosis, thus exerting an anti-heart failure effect^61, 62^.

The novel aspects of our study are as follows. First, we identified RNASE2, MFAP4, CHRDL1, and KCNN3 as potential biomarkers and therapeutic targets for IHF through bioinformatics and three machine learning approaches. Secondly, we validated these four genes in other dataset, and the validation results showed that the diagnostic model composed of these four genes has high diagnostic value, which provides new ideas for our future research on the molecular mechanisms of IHF. In addition, our enrichment analysis and immune infiltration analysis of the dataset showed that the molecular mechanisms of IHF are related to immunity and inflammation, which provides us with ideas for developing new therapeutic modalities for IHF. Nevertheless, there are some shortcomings in this study. Firstly, it is difficult to establish a causal relationship between gene expression differences and the pathophysiological mechanisms of heart failure. Secondly, our dataset was derived from myocardial tissue and the lack of validation of the peripheral blood dataset may limit the application of diagnostic models. Therefore, although our results were validated in other dataset, further clinical trials are needed to confirm our study.

## 5. CONCLUSION

We conducted a bioinformatic analysis of the GEO dataset to investigate the underlying molecular mechanisms of IHF and the immune cell infiltration environment within the failing myocardium. Through the implementation of three machine learning algorithms (LASSO, RF, and SVM-RFE), we have identified RNASE2, MFAP4, CHRDL1, and KCNN3 as potential biomarkers and therapeutic targets for the treatment of IHF. Of particular significance, we have developed diagnostic models and nomogram tools based on these four genes, providing a novel understanding of the pathogenesis of IHF and offering exciting prospects for future in-depth studies.

## Acknowledgements

Not applicable.

## Authors’ contributions

YDY conducted statistical analysis and drafted the article. YDY and YL were involved in the conception and design of the study. YDY contributed to picture processing and article reviewing. YTX reviewed and proofread the article. YTX and YL provided effective scientific suggestions and supervision and created the final revision of the manuscript. All authors read and approved the final manuscript.

## Consent for publication

Not applicable.

## Availability of data and materials

Publicly available datasets were analyzed in this study. This data can be found here: GSE76701; GSE21610; GSE57338.

## Ethics approval and consent to participate

Not applicable.

## REFERENCES

[1] S.A. Hunt, W.T. Abraham, M.H. Chin, A.M. Feldman, G.S. Francis, T.G. Ganiats, M. Jessup, M.A. Konstam, D.M. Mancini, K. Michl, J.A. Oates, P.S. Rahko, M.A. Silver, L.W. Stevenson, C.W. Yancy, E.M. Antman, S.C. Smith Jr., C.D. Adams, J.L. Anderson, D.P. Faxon, V. Fuster, J.L. Halperin, L.F. Hiratzka, S.A. Hunt, A.K. Jacobs, R. Nishimura, J.P. Ornato, R.L. Page, B. Riegel, Acc/aha 2005 guideline update for the diagnosis and management of chronic heart failure in the adult summary article, J. Am. Coll. Cardiol. 46 (2005) 1116–1143, https://doi.org/10.1016/j.jacc2005.08023.

[2] Heidenreich P A, Bozkurt B, Aguilar D, et al. 2022 AHA/ACC/HFSA guideline for the management of heart failure: a report of the American College of Cardiology/American Heart Association Joint Committee on Clinical Practice Guidelines. Journal of the American College of Cardiology, 2022, 79(17): e263-e421.

[3] Peng H, Abdel-Latif A. Cellular Therapy for Ischemic Heart Disease: An Update. Adv Exp Med Biol. 2019;1201:195–213. doi:10.1007/978-3-030-31206-0_10

[4] Kologrivova I, Shtatolkina M, Suslova T, Ryabov V. Cells of the Immune System in Cardiac Remodeling: Main Players in Resolution of Inflammation and Repair After Myocardial Infarction. Front Immunol. 2021;12:664457. Published 2021 Apr 2. doi:10.3389/fimmu.2021.664457

[5] Peet C, Ivetic A, Bromage DI, Shah AM. Cardiac monocytes and macrophages after myocardial infarction. Cardiovasc Res. 2020;116(6):1101–1112. doi:10.1093/cvr/cvz336

[6] Zhang Y, Bauersachs J, Langer HF. Immune mechanisms in heart failure. Eur J Heart Fail. 2017;19(11):1379–1389. doi:10.1002/ejhf.942

[7] Kumar N, Narayan Das N, Gupta D, Gupta K, Bindra J. Efficient Automated Disease Diagnosis Using Machine Learning Models. J Healthc Eng. 2021;2021:9983652. Published 2021 May 4. doi:10.1155/2021/9983652

[8] Schwientek P, Ellinghaus P, Steppan S, et al. Global gene expression analysis in nonfailing and failing myocardium pre-and postpulsatile and nonpulsatile ventricular assist device support. Physiol Genomics. 2010;42(3):397–405. doi:10.1152/physiolgenomics.00030.2010

[9] Kim EH, Galchev VI, Kim JY, et al. Differential protein expression and basal lamina remodeling in human heart failure. Proteomics Clin Appl. 2016;10(5):585–596. doi:10.1002/prca.201500099

[10] Liu Y, Morley M, Brandimarto J, et al. RNA-Seq identifies novel myocardial gene expression signatures of heart failure. Genomics. 2015;105(2):83–89. doi:10.1016/j.ygeno.2014.12.002

[11] Barrett T, Wilhite SE, Ledoux P, et al. NCBI GEO: archive for functional genomics data sets--update. Nucleic Acids Res. 2013;41(Database issue):D991-D995. doi:10.1093/nar/gks1193

[12] Davis S, Meltzer PS. GEOquery: a bridge between the Gene Expression Omnibus (GEO) and BioConductor. Bioinformatics. 2007 Jul 15;23(14):1846–7. doi: 10.1093/bioinformatics/btm254. Epub 2007 May 12. PMID: 17496320.

[13] Leek JT, Johnson WE, Parker HS, Jaffe AE, Storey JD. The sva package for removing batch effects and other unwanted variation in high-throughput experiments. Bioinformatics. 2012 Mar 15;28(6):882–3. doi: 10.1093/bioinformatics/bts034. Epub 2012 Jan 17. PMID: 22257669; PMCID: PMC3307112.

[14] Gu Z, Eils R, Schlesner M. Complex heatmaps reveal patterns and correlations in multidimensional genomic data. Bioinformatics. 2016 Sep 15;32(18):2847–9. doi: 10.1093/bioinformatics/btw313. Epub 2016 May 20. PMID: 27207943.

[15] Smyth G K. Limma: linear models for microarray data. In Bioinformatics and computational biology solutions using R and Bioconductor. 2013.

[16] Langfelder P, Horvath S. WGCNA: an R package for weighted correlation network analysis. BMC Bioinformatics. 2008;9:559. Published 2008 Dec 29. doi:10.1186/1471-2105-9-559

[17] Yu G, Wang LG, Han Y, He QY. clusterProfiler: an R package for comparing biological themes among gene clusters. OMICS. 2012;16(5):284–287. doi:10.1089/omi.2011.0118

[18] The Gene Ontology Consortium. The Gene Ontology Resource: 20 years and still GOing strong. Nucleic Acids Res. 2019;47(D1):D330–D338. doi:10.1093/nar/gky1055

[19] Kanehisa M, Goto S. KEGG: kyoto encyclopedia of genes and genomes. Nucleic Acids Res. 2000;28(1):27–30. doi:10.1093/nar/28.1.27

[20] Subramanian A, Tamayo P, Mootha VK, et al. Gene set enrichment analysis: a knowledge-based approach for interpreting genome-wide expression profiles. Proc Natl Acad Sci U S A. 2005;102(43):15545–15550. doi:10.1073/pnas.0506580102

[21] Szklarczyk D, Gable AL, Nastou KC, et al. The STRING database in 2021: customizable protein-protein networks, and functional characterization of user-uploaded gene/measurement sets [published correction appears in Nucleic Acids Res. 2021 Oct 11;49(18):10800]. Nucleic Acids Res. 2021;49(D1):D605-D612. doi:10.1093/nar/gkaa1074

[22] Shannon P, Markiel A, Ozier O, et al. Cytoscape: a software environment for integrated models of biomolecular interaction networks. Genome Res. 2003;13(11):2498–2504. doi:10.1101/gr.1239303

[23] Friedman J, Hastie T, Tibshirani R. Regularization Paths for Generalized Linear Models via Coordinate Descent. J Stat Softw. 2010;33(1):1–22.

[24] Petralia F, Wang P, Yang J, Tu Z. Integrative random forest for gene regulatory network inference. Bioinformatics. 2015;31(12):i197–i205. doi:10.1093/bioinformatics/btv268

[25] Huang S, Cai N, Pacheco PP, Narrandes S, Wang Y, Xu W. Applications of Support Vector Machine (SVM) Learning in Cancer Genomics. Cancer Genomics Proteomics. 2018;15(1):41–51. doi:10.21873/cgp.20063

[26] Gou M, Qian N, Zhang Y, et al. Construction of a nomogram to predict the survival of metastatic gastric cancer patients that received immunotherapy. Front Immunol. 2022;13:950868. Published 2022 Sep 26. doi:10.3389/fimmu.2022.950868

[27] Ye L, Zhang T, Kang Z, et al. Tumor-Infiltrating Immune Cells Act as a Marker for Prognosis in Colorectal Cancer. Front Immunol. 2019;10:2368. Published 2019 Oct 17. doi:10.3389/fimmu.2019.02368

[28] Newman AM, Liu CL, Green MR, et al. Robust enumeration of cell subsets from tissue expression profiles. Nat Methods. 2015;12(5):453–457. doi:10.1038/nmeth.3337

[29] Del Buono MG, Moroni F, Montone RA, Azzalini L, Sanna T, Abbate A. Ischemic Cardiomyopathy and Heart Failure After Acute Myocardial Infarction. Curr Cardiol Rep. 2022;24(10):1505–1515. doi:10.1007/s11886-022-01766-6

[30] Ngwenyama N, Kaur K, Bugg D, et al. Antigen presentation by cardiac fibroblasts promotes cardiac dysfunction. Nat Cardiovasc Res. 2022;1(8):761–774. doi:10.1038/s44161-022-00116-7

[31] Kumar V, Prabhu SD, Bansal SS. CD4+ T-lymphocytes exhibit biphasic kinetics post-myocardial infarction. Front Cardiovasc Med. 2022;9:992653. Published 2022 Aug 25. doi:10.3389/fcvm.2022.992653

[32] Gage JR, Fonarow G, Hamilton M, Widawski M, Martínez-Maza O, Vredevoe DL. Beta blocker and angiotensin-converting enzyme inhibitor therapy is associated with decreased Th1/Th2 cytokine ratios and inflammatory cytokine production in patients with chronic heart failure. Neuroimmunomodulation. 2004;11(3):173–180. doi:10.1159/000076766

[33] Bi HL, Zhang XL, Zhang YL, et al. The deubiquitinase UCHL1 regulates cardiac hypertrophy by stabilizing epidermal growth factor receptor. Sci Adv. 2020;6(16):eaax4826. Published 2020 Apr 17. doi:10.1126/sciadv.aax4826

[34] Nishida K, Otsu K. Sterile Inflammation and Degradation Systems in Heart Failure. Circ J. 2017;81(5):622–628. doi:10.1253/circj.CJ-17-0261

[35] Brown DI, Parry TL, Willis MS. Ubiquitin Ligases and Posttranslational Regulation of Energy in the Heart: The Hand that Feeds. Compr Physiol. 2017;7(3):841–862. Published 2017 Jun 18. doi:10.1002/cphy.c160024

[36] Pokorna Z, Jirkovsky E, Hlavackova M, et al. In vitro and in vivo investigation of cardiotoxicity associated with anticancer proteasome inhibitors and their combination with anthracycline. Clin Sci (Lond). 2019;133(16):1827–1844. Published 2019 Aug 27. doi:10.1042/CS20190139

[37] Gupta SK, Haigh BJ, Griffin FJ, Wheeler TT. The mammalian secreted RNases: mechanisms of action in host defence. Innate Immun. 2013;19(1):86–97. doi:10.1177/1753425912446955

[38] Bedoya VI, Boasso A, Hardy AW, Rybak S, Shearer GM, Rugeles MT. Ribonucleases in HIV type 1 inhibition: effect of recombinant RNases on infection of primary T cells and immune activation-induced RNase gene and protein expression. AIDS Res Hum Retroviruses. 2006;22(9):897–907. doi:10.1089/aid.2006.22.897

[39] Yang D, Chen Q, Rosenberg HF, et al. Human ribonuclease A superfamily members, eosinophil-derived neurotoxin and pancreatic ribonuclease, induce dendritic cell maturation and activation. J Immunol. 2004;173(10):6134–6142. doi:10.4049/jimmunol.173.10.6134

[40] Yang D, Chen Q, Su SB, et al. Eosinophil-derived neurotoxin acts as an alarmin to activate the TLR2-MyD88 signal pathway in dendritic cells and enhances Th2 immune responses. J Exp Med. 2008;205(1):79–90. doi:10.1084/jem.20062027

[41] Yang D, Rosenberg HF, Chen Q, Dyer KD, Kurosaka K, Oppenheim JJ. Eosinophil-derived neurotoxin (EDN), an antimicrobial protein with chemotactic activities for dendritic cells. Blood. 2003;102(9):3396–3403. doi:10.1182/blood-2003-01-0151

[42] Ostendorf T, Zillinger T, Andryka K, et al. Immune Sensing of Synthetic, Bacterial, and Protozoan RNA by Toll-like Receptor 8 Requires Coordinated Processing by RNase T2 and RNase 2. Immunity. 2020;52(4):591-605.e6. doi:10.1016/j.immuni.2020.03.009

[43] Zhang XY, Huang Z, Li QJ, et al. Ischemic postconditioning attenuates the inflammatory response in ischemia/reperfusion myocardium by upregulating miR-499 and inhibiting TLR2 activation. Mol Med Rep. 2020;22(1):209–218. doi:10.3892/mmr.2020.11104

[44] Ong SLM, de Vos IJHM, Meroshini M, Poobalan Y, Dunn NR. Microfibril-associated glycoprotein 4 (Mfap4) regulates haematopoiesis in zebrafish. Sci Rep. 2020;10(1):11801. Published 2020 Jul 16. doi:10.1038/s41598-020-68792-8

[45] Schlosser A, Pilecki B, Hemstra LE, et al. MFAP4 Promotes Vascular Smooth Muscle Migration, Proliferation and Accelerates Neointima Formation. Arterioscler Thromb Vasc Biol. 2016;36(1):122–133. doi:10.1161/ATVBAHA.115.306672

[46] Wulf-Johansson H, Lock Johansson S, Schlosser A, et al. Localization of microfibrillar-associated protein 4 (MFAP4) in human tissues: clinical evaluation of serum MFAP4 and its association with various cardiovascular conditions. PLoS One. 2013;8(12):e82243. Published 2013 Dec 13. doi:10.1371/journal.pone.0082243

[47] Han C, Peng Y, Yang X, et al. Declined plasma microfibrillar-associated protein 4 levels in acute coronary syndrome. Eur J Med Res. 2023;28(1):32. Published 2023 Jan 18. doi:10.1186/s40001-023-01002-z

[48] Kanaan R, Medlej-Hashim M, Jounblat R, Pilecki B, Sorensen GL. Microfibrillar-associated protein 4 in health and disease. Matrix Biol. 2022;111:1–25. doi:10.1016/j.matbio.2022.05.008

[49] Dorn LE, Lawrence W, Petrosino JM, et al. Microfibrillar-Associated Protein 4 Regulates Stress-Induced Cardiac Remodeling. Circ Res. 2021;128(6):723–737. doi:10.1161/CIRCRESAHA.120.317146

[50] Pei YF, Zhang YJ, Lei Y, Wu WD, Ma TH, Liu XQ. Hypermethylation of the CHRDL1 promoter induces proliferation and metastasis by activating Akt and Erk in gastric cancer [published correction appears in Oncotarget. 2017 Jul 18;8(29):48526. Wu, Ding-Wei [corrected to Wu, Wei-ding]]. Oncotarget. 2017;8(14):23155-23166. doi:10.18632/oncotarget.15513

[51] Ruozi G, Bortolotti F, Mura A, et al. Cardioprotective factors against myocardial infarction selected in vivo from an AAV secretome library. Sci Transl Med. 2022;14(660):eabo0699. doi:10.1126/scitranslmed.abo0699

[52] Natarajan P, Pampana A, Graham SE, et al. Chromosome Xq23 is associated with lower atherogenic lipid concentrations and favorable cardiometabolic indices. Nat Commun. 2021;12(1):2182. Published 2021 Apr 12. doi:10.1038/s41467-021-22339-1

[53] National Center for Biotechnology Information. PubChem Gene Summary for Gene 3782. https://pubchem.ncbi.nlm.nih.gov/gene/KCNN3/human. Accessed Mar. 29, 2023.

[54] Ellinor PT, Lunetta KL, Glazer NL, et al. Common variants in KCNN3 are associated with lone atrial fibrillation. Nat Genet. 2010;42(3):240–244. doi:10.1038/ng.537

[55] Darkow E, Nguyen TT, Stolina M, et al. Small Conductance Ca2 +-Activated K+ (SK) Channel mRNA Expression in Human Atrial and Ventricular Tissue: Comparison Between Donor, Atrial Fibrillation and Heart Failure Tissue. Front Physiol. 2021;12:650964. Published 2021 Apr 1. doi:10.3389/fphys.2021.650964

[56] Ortega A, Tarazón E, Roselló-Lletí E, et al. Patients with Dilated Cardiomyopathy and Sustained Monomorphic Ventricular Tachycardia Show Up-Regulation of KCNN3 and KCNJ2 Genes and CACNG8-Linked Left Ventricular Dysfunction. PLoS One. 2015;10(12):e0145518. Published 2015 Dec 28. doi:10.1371/journal.pone.0145518

[57] Mouton AJ, Li X, Hall ME, Hall JE. Obesity, Hypertension, and Cardiac Dysfunction: Novel Roles of Immunometabolism in Macrophage Activation and Inflammation. Circ Res. 2020;126(6):789–806. doi:10.1161/CIRCRESAHA.119.312321

[58] Zhang L, Chen J, Yan L, He Q, Xie H, Chen M. Resveratrol Ameliorates Cardiac Remodeling in a Murine Model of Heart Failure With Preserved Ejection Fraction [published correction appears in Front Pharmacol. 2022 Mar 18;13:857367]. Front Pharmacol. 2021;12:646240. Published 2021 Jun 10. doi:10.3389/fphar.2021.646240

[59] Balligand JL, Ungureanu D, Kelly RA, et al. Abnormal contractile function due to induction of nitric oxide synthesis in rat cardiac myocytes follows exposure to activated macrophage-conditioned medium. J Clin Invest. 1993;91(5):2314–2319. doi:10.1172/JCI116461

[60] Wrigley BJ, Lip GY, Shantsila E. The role of monocytes and inflammation in the pathophysiology of heart failure. Eur J Heart Fail. 2011;13(11):1161–1171. doi:10.1093/eurjhf/hfr122

[61] Blanco-Domínguez R, de la Fuente H, Rodríguez C, et al. CD69 expression on regulatory T cells protects from immune damage after myocardial infarction. J Clin Invest. 2022;132(21):e152418. Published 2022 Nov 1. doi:10.1172/JCI152418

[62] Yan X, Shichita T, Katsumata Y, et al. Deleterious effect of the IL-23/IL-17A axis and γδT cells on left ventricular remodeling after myocardial infarction. J Am Heart Assoc. 2012;1(5):e004408. doi:10.1161/JAHA.112.004408

